# Asymmetrical representation of the upper and lower visual fields in oculomotor maps

**DOI:** 10.1101/086397

**Authors:** Zhiguo Wang, Benchi Wang, Matthew Finkbeiner

**Affiliations:** Institute of Psychological Sciences, Hangzhou Normal University, Hangzhou, China; Center for Cognition and Brain Disorders, Hangzhou Normal University, Hangzhou, China; Department of Cognitive Psychology, Vrije Universiteit Amsterdam, Amsterdam, the Netherlands; Department of Cognitive Science, Macquarie University, Sydney, Australia

## Abstract

The striate area devoted to the lower visual field (LVF) is larger than that devoted to the upper visual field (UVF). A similar anatomical asymmetry also exists in the LGN. Here we take advantage of two experimental tasks that are known to modulate the direction and amplitude of saccades to demonstrate a visual field asymmetry in oculomotor maps. Participants made visually guided saccades. In Experiment 1, the saccade target was accompanied by a visual distractor. The distractor’s presence modulated the direction of saccades, and this effect was much stronger for LVF targets. In Experiment 2, the temporal gap between the offset of the fixation stimulus and the onset of the saccade target was manipulated. This manipulation modulated the amplitude of saccades and this modulation was stronger for saccades towards UVF targets. Taken together, these results suggest that the representation of both meridians and eccentricities in the LVF is compressed in oculomotor maps.

## Introduction

Many vision-related brain structures, such as the striate cortex (Sereno et al., 1995; Silver & Kastner, 2009; Van Essen, Newsome, & Maunsell, 1984; Wandell, Dumoulin, & Brewer, 2007), the lateral geniculate nucleus (LGN; Connolly & Van Essen, 1984), and the superior colliculus (SC; Robinson, 1972; Schneider & Kastner, 2005), are organized into gaze-centered, or retinotopic, maps. The mapping between the visual field and these maps has a characterizing central-peripheral asymmetry—the central visual field is represented by disproportionally large brain areas, whereas more eccentric regions of the visual field are represented by increasingly smaller brain areas. This anatomical asymmetry seems to reflect the photoreceptor density on the retina and the visual processing efficiency of the central and peripheral vision. Most notably, visual acuity exponentially decreases as the eccentricity increases (e.g., Duncan & Boynton, 2003). In addition to this central-peripheral asymmetry, there is also a functional asymmetry between visual fields above and below fixation. Previous studies have shown various processing advantages in the lower visual fields (LVF; for a review, see Skrandies, 1987). For instance, attention has been shown to have finer resolution in the LVF (He, Cavanagh, & Intriligator, 1996), and vision is more sensitive in the LVF (e.g., Carrasco, Talgar, & Cameron, 2001; Talgar & Carrasco, 2002). This functional asymmetry may also have its neuroanatomical basis. For instance, more cones on the retina are dedicated to processing LVF visual information; in non-human primates, the neural tissues representing the LVF is about 1.2 times of that representing the UVF in LGN (Connolly & Van Essen, 1984) and in the striate cortex (Tootell, Switkes, Silverman, & Hamilton, 1988; Van Essen et al., 1984). Human imaging studies have also shown that visual stimuli in the LVF evoke stronger (and possibly also larger) neural activation in the striate cortex (Liu, Heeger, & Carrasco, 2006). Together, these findings seem to suggest that functional asymmetries that exist in the visual field all have their anatomical basis.

One often overlooked functional asymmetry of the UVF and LVF exists in the oculomotor system. It has been long known that saccadic eye movements, or saccades, to the LVF takes longer to program (Heywood & Churcher, 1980), an observation that has been recently demonstrated in a variety of experimental conditions (Abegg, Pianezzi, & Barton, 2015). Similar asymmetries also exist in saccade amplitudes and directions (Petrova & Wentura, 2012; Wang & Theeuwes, 2014). These asymmetries are not directly associated with visual perception and may arise from neural structures responsible for saccade programming. According to Previc (1990), evolution has led to a functional specialization of the UVF and LVF. The LVF is more relevant to visual processing in “near” space whereas the UVF is specialized for processing in “far” space. The processing of visual information in far space frequently involves the shift of gaze. Since previously observed perceptual advantages in the LVF are likely linked to an overrepresentation of the LVF in brain regions responsible for vision processing (e.g., V1), it seems probable that the UVF advantages in oculomotor behaviors are similarly linked to an overrepresentation of the UVF in oculomotor maps, such as the superior colliculus (SC) and the frontal eye fields (FEFs).

Both the SC and the FEFs contain topographical maps that encodes the direction and amplitude of saccades. The topographical organization of the FEF is rather coarse (Sommer & Wurtz, 2000) while that of the SC is much more fine-grained, similar to that of V1 (Schwartz, 1977). Our current knowledge about the shape of the SC motor map was largely based on the findings of Robinson (1972), in which micro-stimulation was delivered to various SC sites to evoke saccades of varying directions and amplitudes. Based on 50 SC stimulation sites from two monkeys, Robinson (1972) constructed a contour map representing equal amplitudes and directions (see Figure 1). This map assumed symmetrical mapping of the UVF and LVF in the SC. However, it is important to note that this map is constructed by first dividing the stimulation sites into strips of constant amplitude and direction “*by eye*” (see Robinson, 1972, pp. 1800-1801); it was never intended to be regarded as a *precise* description of the SC motor map, but rather a close approximation. There is no evidence, so far, to convincingly suggest that the LVF and UVF are mapped symmetrically in the SC and other oculomotor maps. Here we take advantage of two well established experimental tasks that are known to modulate the direction and amplitude of saccades (Wang & Theeuwes, 2014) to show that the LVF is underrepresented in oculomotor maps, presumably in the SC (and possibly also in the FEFs).

**Figure 1.**
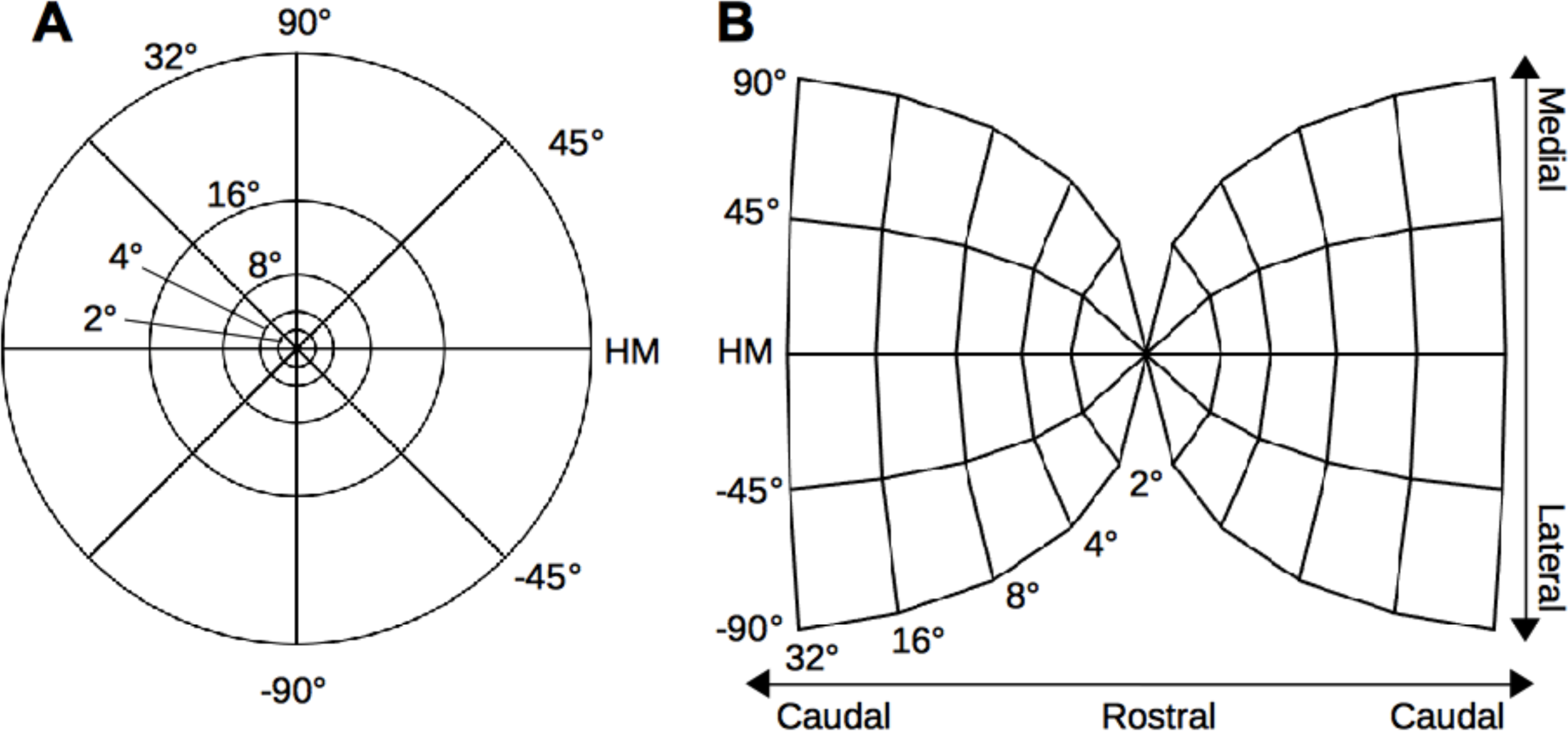
(A) The visual space represented in polar coordinates. (B) The SC motor map depicted by the mapping functions proposed by Ottes, van Gisbergen and Eggermont (1986).

In two experiments, participants made saccades to an onset visual target. In Experiment 1, the saccade target was accompanied by a visual distractor (see Figure 2A). The visual distractor has been shown to either attract or repel the direction of saccades, depending on its angular distance to the saccade target (e.g., McSorley, Cruickshank, & Inman, 2009; Wang & Theeuwes, 2014). In Experiment 2, the fixation stimulus was removed either before or after the presentation of the saccade target (see Figure 2B). This manipulation is known to influence saccade latencies (Reuter-Lorenz, Hughes, & Fendrich, 1991; Saslow, 1967; Walker, Deubel, Schneider, & Findlay, 1997), and more importantly, saccade amplitudes (Wang & Theeuwes, 2014). To anticipate our results, we find a robust effect of the distractor’s presence on saccade directions as well as a robust effect of the temporal gap’s presence on saccade amplitudes. Critically, both of these effects were asymmetrical, with stronger distractor effects on saccade direction in the LVF and stronger temporal gap effects on saccade amplitude in the UVF. We will argue that, taken together, these findings point to an underrepresentation of the LVF in oculomotor maps, likely in the SC and the FEFs.

**Figure 2.**
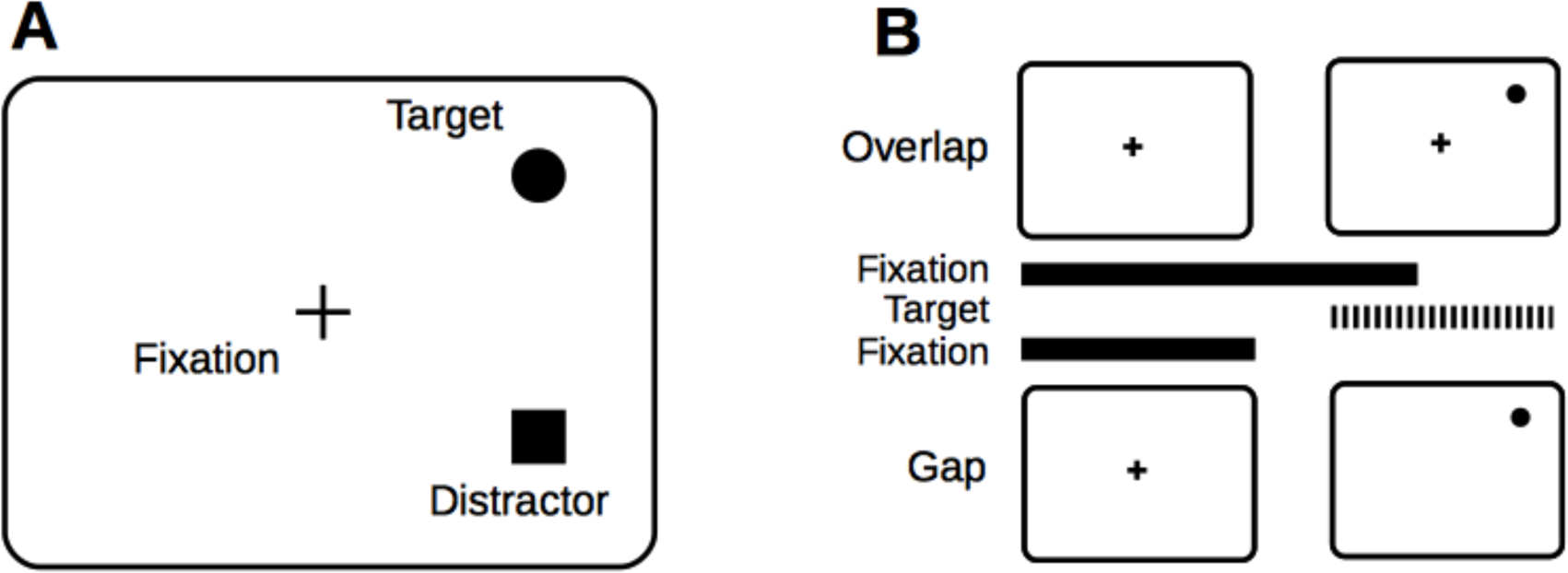
The distractor task (A) and the temporal gap/overlap task (B). The presence of visual distractors has been shown to push the saccade direction in opposite direction, when its angular distance to the target is over ~30º. In the temporal gap/overlap task, the fixation stimulus (cross in the illustration) is extinguished before or after the onset of the saccade target. The temporal gap has been shown to reduce both saccade latency and amplitudes.

## Experiment 1: Saccade direction

With a distractor task, Experiment 1 aims to examine whether meridians in the UVF and LVF are symmetrically represented in oculomotor maps. To this end, the target and distractor were presented at locations mirroring the horizontal meridian. That is, if the saccade target were presented at 30° (polar angle) above the horizontal meridian, the distractor would then be presented 30° below. The saccade targets were presented randomly in either the UVF or LVF (equal probability). The rationale was straightforward. It has been shown that, all other things being equal, the directional deviation caused by a visual distractor is determined by its collicular distance from the saccade target (Wang, Kruijne, & Theeuwes, 2012; Wang & Theeuwes, 2014). In our experiment, the collicular distance between the target and the distractor is the same regardless of whether the target appears in the UVF or LVF. If we assume a symmetrical representation of the UVF and LVF in the SC, the directional deviation in the observed saccades that is caused by the distractor should be comparable in magnitude across the visual fields. In contrast, if the LVF is relatively compressed in the SC motor map (as illustrated in Figure 3A), though the magnitude of deviation *in collicular space* (marked by arrows) is comparable across visual fields, the collicular deviation for LVF targets will map to a much larger directional change *in visual space*.

**Figure 3.**
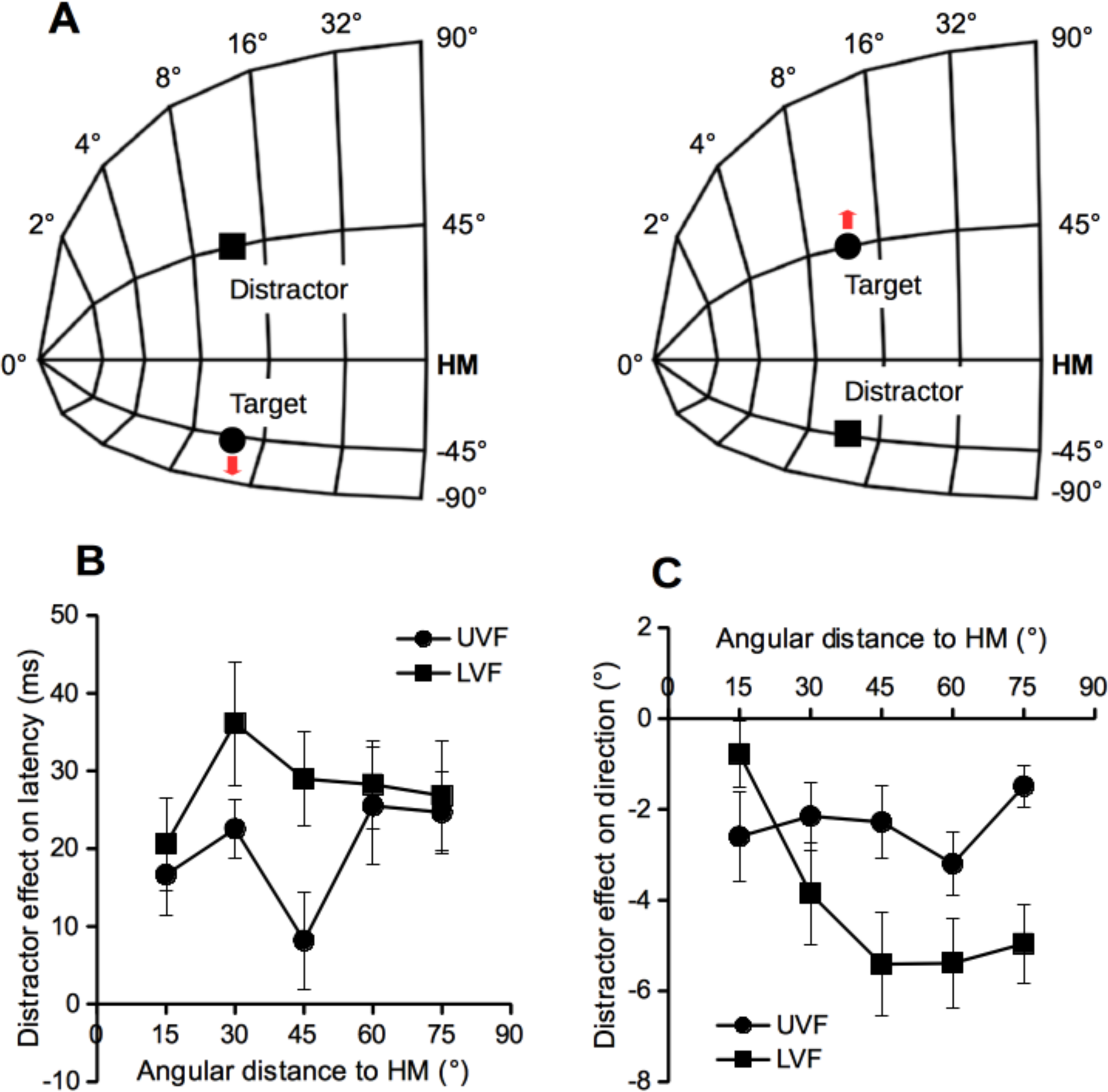
(A) Compressed representation of the LVF in the SC will lead to stronger deviation of saccade direction. When the target and distractor mirror the horizontal meridian, the same amount of collicular deviation (red arrow) will be produced regardless whether the target is in the UVF or LVF. However, because the LVF is compressed in the SC, the same amount of collicular deviation actually maps to a much larger change in direction for saccades to LVF targets. (B-C) The effect of distractors on saccade latency and initial saccade direction. HM, horizontal meridian; UVF, upper visual field; LVF, lower visual field. Error bars denote ±1 SEM.

Preliminary results of this experiment was briefly presented at the 2014 Vision Sciences Society Meeting (Wang, 2014).

## Method

The research protocols reported here were approved by the Institutional Review Board of Center for Cognition and Brain Disorders, Hangzhou Normal University. All participants gave written informed consent.

### Participants

Experiment 1 had 19 participants, one participant (6_XJY) was excluded because she did not complete the task; another participant (1_XSX) was also excluded because the eye movement data file was corrupted due to device malfunction. The remaining 17 participants (7 males, mean age: 23.41 years) all had normal or corrected-to-normal vision, and reported no history of psychological or neurological disorders. They were paid 40 *yuan* per hour for their participation.

### Apparatus

The experiment was carried out in a dimly lit laboratory. All stimuli were presented against a black background (1.5 cd/m^2^) on a 17-inch CRT monitor. The visible area of the monitor measured 32° × 24° visual angle at a view distance of ~71 cm (maintained by using a chinrest). The participant’s gaze direction was monitored and recorded with an EyeLink 1000 (SR Research) eye tracker, sampling at 500 Hz or 1000 Hz. The spatial resolution of this eye tracker was 0.2° or better.

### Task procedure

A trial started with the presentation of a gray fixation cross that measured 1° × 1° (Weber contrast = 8.69). After a delay of 500-700 ms (randomly selected), a saccade target (grey circle; diameter = 1°, Weber contrast = 8.69) appeared at an eccentricity of 7.7°. A visual distractor (filled gray square; 1° x 1°, Weber contrast = 8.69) appeared at the same time on 50% of the trials (see Figure 2A). To discourage anticipatory response, the target was not presented on 1/16 of the trials, regardless of the presentation of the distractor. The participant had 1000 ms to initiate a saccade to the target if it appeared on display. After a random inter-trial interval of 750-1500 ms, the next trial began.

The target and distractor could deviate from the horizontal meridian for 15°, 30°, 45°, 60°, or 75° (polar angle). When presented together, they always appeared at locations mirroring the horizontal meridian. Thus, the angular separation between them could be 30°, 60°, 90°, 120° or 150°. By design, a total of 640 trials were tested. A block of 24 practice trials was provided before formal testing. During testing, the participants could take a break after every 160 trials or whenever they felt need it. The eye tracker was calibrated at the beginning of the testing session and after each break. Warning messages were displayed and a trial was flagged as an “error trial” if the participants failed to maintain fixation before target onset, failed to respond within 1000 ms, or the primary saccade missed the saccade target by more than 2°. All error trials were later presented to the participants in a random order, until all trials were successfully completed.

### Dependent measures

The dependent measures of prime interest were saccade latencies and directional deviations caused by the distractor. Saccade latency was the time interval between the onset of the stimulus (target, distractor, or both) and the onset of the primary saccade. Saccade onset was detected online with a velocity threshold of 30°/s and an acceleration threshold of 8000°/s^2^. The initial direction of the primary saccade was determined with its start position and the gaze position at which the saccade had traversed 30% of its duration (for similar measures, see Van der Stigchel, Meeter, & Theeuwes, 2007; Wang & Theeuwes, 2014). Directional deviation was quantified as the difference in initial saccade direction between distractor-present and distractor-absent trials, with positive and negative values denoting deviation towards and away from the distractors, respectively.

## Results

Only successfully completed trials were analyzed. Trials were excluded if they met one of the following criteria: a) the latency of the primary saccade was below 80 ms or above 550 ms (1.53%), b) the duration of the primary saccade was longer than 100 ms (2.38%), c) the start position of the primary saccade deviated more than 2° from central fixation (3.49%), or d) the initial direction of the primary saccade deviated from the target direction for more than 45° (3.53%). After data cleaning, 91.18% of the trials remained.

### Saccade latency

Mean saccade latencies are presented in Table 1. A repeated measures ANOVA revealed significant main effects for distractor presence (with vs. without distractor), *F*(1,16) = 44.13, *p* < 0.001, visual field (target in UVF vs. LVF), *F*(1,16) = 50.62, *p* < 0.001, and distance to horizontal meridian (in polar angle), *F*(4,64) = 20.38, *p* < 0.001. These effects occurred because saccade latencies were longer when the distractor was present, longer for LVF targets, and generally increased with distance to horizontal meridian. A significant two-way interaction occurred between visual field and distance to horizontal meridian, *F*(4,64) = 8.00, *p* < 0.001; saccade latencies increased with the distance to horizontal meridian and the slope was much steeper for targets in the LVF (see Table 1). A marginal two-way interaction occurred between visual field and distractor presence, *F*(4,64) = 4.40, *p* = 0.05; the distractor effect on saccade latency was generally larger for LVF targets (see Figure 3B). All other effects were not significant, all *F* < 1.82, all *p*> 0.16.

**Table 1.**
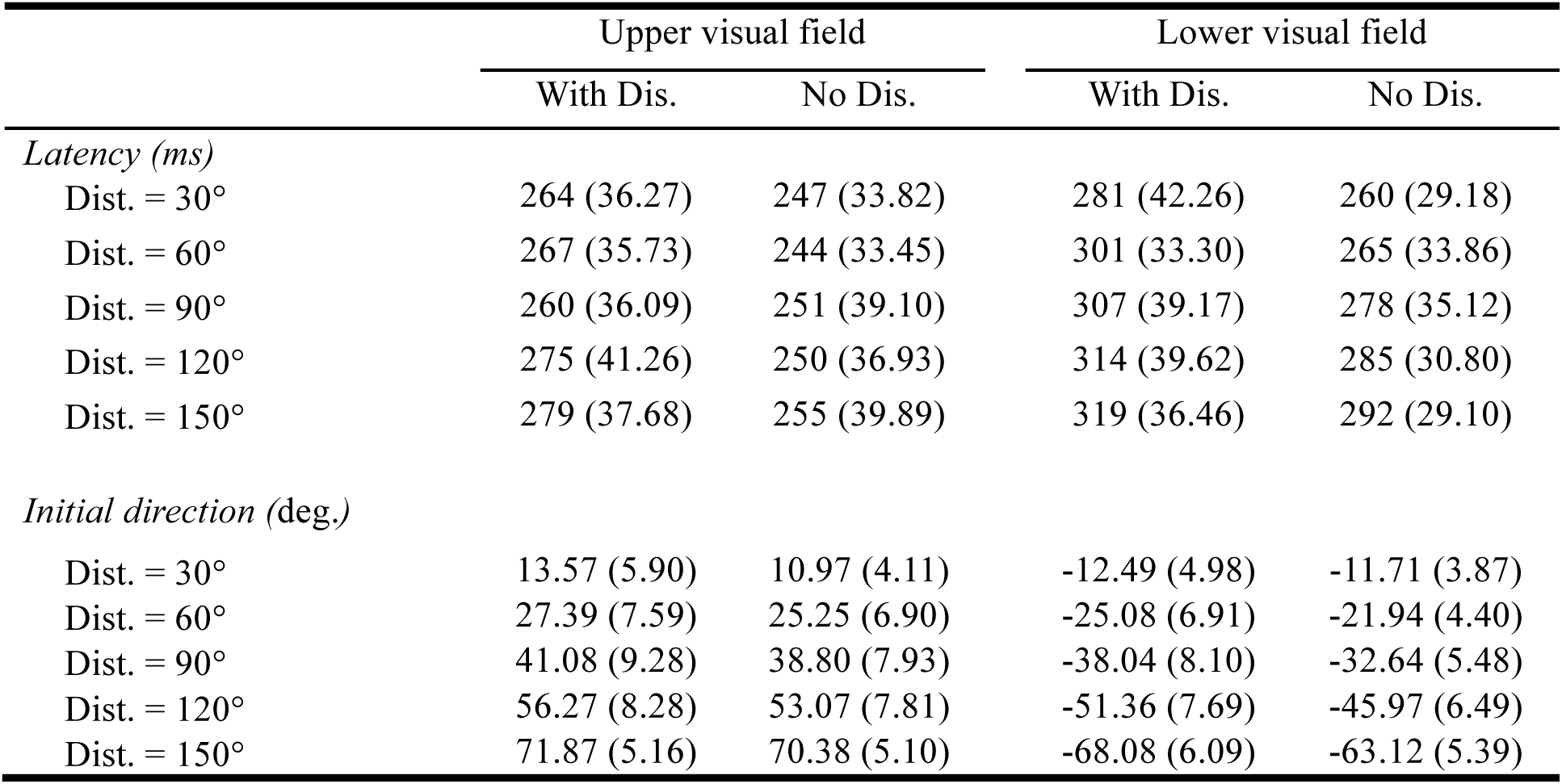
Mean saccade latency and initial saccade direction of each condition in Experiment 1. Numbers in the parentheses are standard deviations (SDs).

### Directional deviation

Directional deviations are presented in Figure 3C. An ANOVA revealed significant main effects for visual field, *F*(1, 16) = 10.87, *p* < 0.01, and distance to horizontal meridian, *F*(4, 64) = 5.29, *p* < 0.001. The directional deviation increased with the distance to horizontal meridian and was much stronger for LVF targets. Importantly, the two-way interaction between visual field and distance to horizontal meridian was significant, *F*(4,64) = 6.24, *p* <0.001. Paired t-tests revealed stronger directional deviations for saccades towards LVF targets when the target was 30° and further away from the horizontal meridian, all *t* > 2.06, all *p* <0.025. When the target was 15° away from the horizontal meridian, the directional deviation was weaker for saccades directed to LVF targets, *t*(16) = 1.95, *p* = 0.035 (1-tailed).

## Discussion

As clearly shown in Figure 3C, the presence of visual distractors pushed the direction of saccades away. In this experiment the target and distractor always appeared at locations mirroring the horizontal meridian. If the UVF and LVF were symmetrically represented in oculomotor maps, then the collicular size of the target and distractor, and the collicular distance between them should be the same regardless of whether the target appeared in the UVF or LVF. Consequently, the directional deviation caused by the distractor should be comparable between targets in the UVF and LVF. But this is not what we found. When targets were relatively far from the horizontal meridian, we found much stronger directional deviation for targets in the LVF. One straight forward, albeit speculative, account is that, as schematically illustrated in Figure 3A, the SC and other oculomotor maps use less neural tissue to represent the LVF. If we assume on this account that (a) a distractor at a fixed distance from the target produces the same amount of deviation in collicular space regardless of whether the target is in the UVF or LVF, and (b) that the LVF is compressed in the SC, then it would follow that the distractor-induced deviations in collicular space would produce a much larger directional change in saccades to LVF targets. This is precisely what we observed. Additionally, as clearly shown in Figure 3C, the difference in directional deviation between the LVF and UVF was more prominent when the targets were 30° or further away from the horizontal meridian, suggesting that the compression of visual field is more obvious in LVF space close to the vertical meridian.

We would like to note that the present observation cannot be explained in terms of perceptual advantages in LVF (e.g., Carrasco et al., 2001; Talgar & Carrasco, 2002). First, the perceptual advantage observed in LVF is largely restricted on the vertical meridian (e.g., Cameron, Tai, & Carrasco, 2002; Talgar & Carrasco, 2002), whereas in the present experiment the target and distractor were never presented on the vertical meridian. Second, the LVF perceptual advantage on the vertical meridian is most noticeable for medium and high frequency stimuli (Cameron et al., 2002; Skrandies, 1987). Even though our saccade target may appear at locations close to the vertical meridian, it was unlikely that this LVF advantage would have affected the perceptual processing of them, which had very low spatial frequency. Finally and most importantly, none of the LVF perceptual advantage findings could explain why response times (saccade latency) were longer for LVF targets.

Previous studies have found that the directional deviation caused by visual distractors increases with saccade latency, possibly because the distractor has more time to compete with the target for selection when saccade latencies are longer (e.g., McSorley, Haggard, & Walker, 2006; Theeuwes & Godijn, 2004). Following from this, it would be reasonable to think that the stronger directional deviation for LVF targets observed in the present experiment might be due to the longer saccade latencies for LVF targets. To examine this possibility, we followed previous work (e.g., McSorley et al., 2009, 2006; Theeuwes & Godijn, 2004) by binning the saccade latencies separately for UVF and LVF targets and then created delta plots for the different conditions. As is clear from Figure 4, while there is a general trend for directional deviations to increase with saccade latency, the difference in directional deviations across visual fields persisted even when the saccade latencies were comparable.

**Figure 4.**
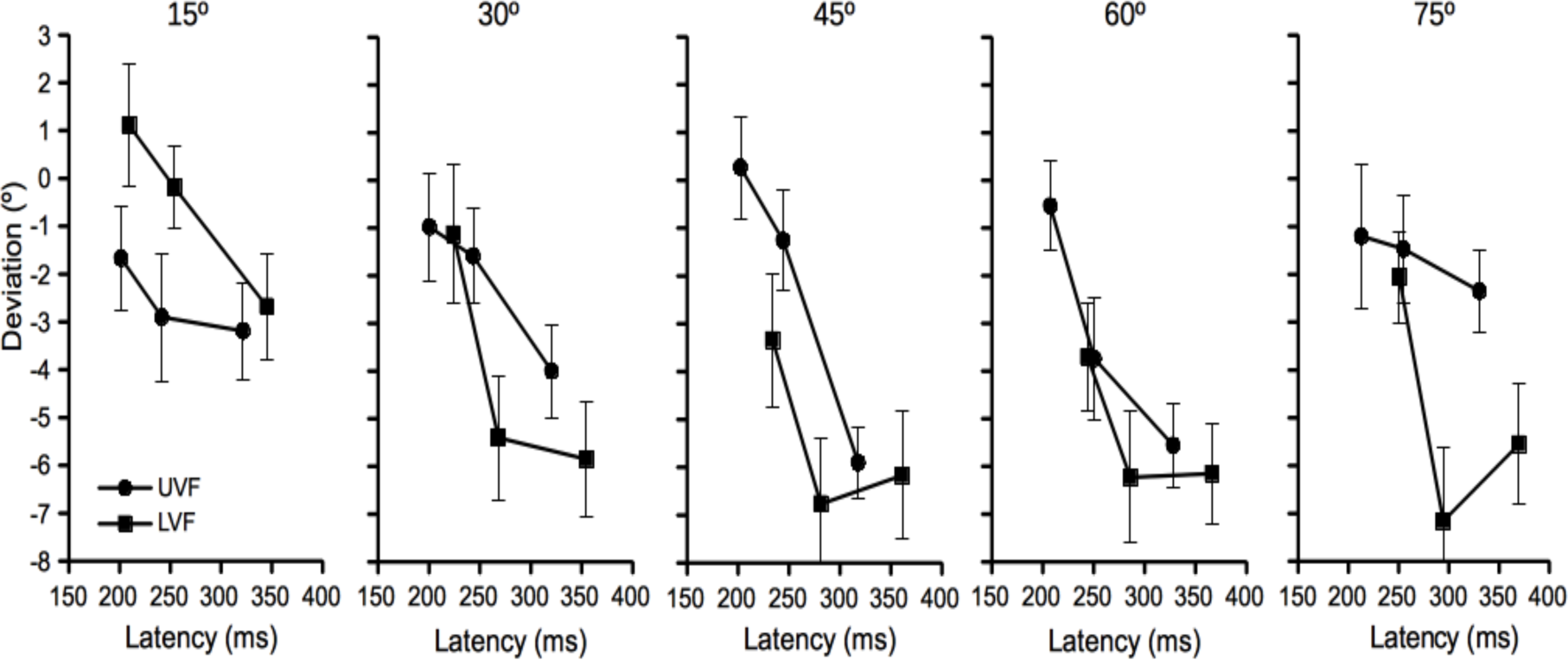
The effect of saccade latency on directional deviation. In this figure, trials were binned according to the saccade latencies. Error bars denote ±1 SEM.

## Experiment 2: Saccade amplitude

The temporal gap between the offset of fixation stimulus and the onset of saccade target has been shown to reduce saccade latencies (e.g., Reuter-Lorenz et al., 1991; Saslow, 1967). A recent study showed that this manipulation also reduces the amplitude of saccades (Wang & Theeuwes, 2014). Dorris and Munoz (1995) found that the temporal gap markedly reduced the activation of SC neurons responsible for maintaining fixation. These fixation neurons are located at the rostral pole of the SC and they globally suppress saccade related activation in more caudal SC regions. Saccade amplitudes are encoded along the rostral-caudal direction; strong fixation activation will repel the saccade-related activation to more caudal SC regions, leading to an increase in saccade amplitude (Wang & Theeuwes, 2014). Experiment 2 was designed to examine if eccentricities in the LVF and UVF are symmetrically encoded in the SC motor map. If so, caudal SC activation evoked by saccade targets at the same eccentricity should receive the same strength of suppression from the rostral pole of the SC. Consequently, a manipulation of temporal gap should produce the same amount of change in saccade amplitudes for LVF and UVF targets. If, though, the representation of eccentricities is compressed in the LVF, then LVF targets will be closer to the fixation in collicular space. As a result, the effect of temporal gap on saccade amplitude should differ across visual fields. This is what we observe in Experiment 2.

## Method

### Participants

This experiment had four participants (all males, mean age: 24.0 years). They were paid 40 *yuan* per hour for their participation.

### Apparatus and stimuli

The experimental setup was the same as Experiment 1, except that the viewing distance was reduced to 61 cm. The saccade target was a white filled disk (diameter = 1°, Weber contrast = 20.5) presented against a black background (1.5 cd/m^2^). The target was presented at an eccentricity of 3°, 6° or 12°, and its angular distance to the horizontal meridian was 30° or 75°.

### Procedure and design

Trials started with the presentation of a white fixation cross (1° × 1°) for 850-1150 ms. The fixation cross was removed either 150 ms before (gap) or 150 ms after (overlap) the onset of the saccade target (see Figure 2B). Longer saccade latencies and larger saccade amplitudes were expected when the fixation stimulus and the saccade target overlapped in time. The participant had 1000 ms to initiate a saccade to the target. After an inter-trial interval of 1000-1500 ms, the next trial began.

Each participant was tested for 5 sessions of 480 trials each. A block of 24 practice trials was provided at the beginning of the first session. Two participants made saccades to targets presented in the left visual field while the other two to targets in the right visual field. A trial was flagged as an “error trial” and later retested if fixation was broken before saccade target onset, or the primary saccade was made to the wrong direction.

## Results

Only successfully completed trials were considered. Those with saccades latencies shorter than 80 ms or longer than 550 ms (2.54%) were excluded from analyses.

### Saccade latency

Mean saccade latencies are presented in Table 2. A repeated measures ANOVA revealed significant main effects for temporal gap (gap vs. overlap), *F*(1,3) = 126.2, *p* < 0.01, angular distance to horizontal meridian (30° vs. 75°), *F*(1,3) = 68.32, *p* < 0.01, and visual field (target in UVF vs. LVF), *F*(1,3) = 16.1, *p* < 0.05. These effects occurred because saccade latencies were longer in the overlap condition, longer for LVF targets, and longer for targets that were further away from the horizontal meridian. The main effect of target eccentricity (3°, 6°, vs. 12°) approached significance, *F*(2,6) = 3.57, *p* = 0.095, and there was trend for latency to increase with target eccentricity. Two-way interactions occurred between temporal gap and target eccentricity, *F*(2,6) = 20.43, *p* < 0.01, because the latency difference between overlap and gap conditions increased with target eccentricity, and between temporal gap and visual field, *F*(1,3) = 37.12, *p* < 0.01, because the temporal gap effect was stronger in the LVF, and between angular distance to horizontal meridian and visual field, *F*(1, 3) = 10.2, *p* < 0.05, because the latency difference between the UVF and LVF was more prominent when the target was 75° away from the horizontal meridian (see Figure 5A-B). All other interactions were not significant, all *F* < 4.38, all *p*> 0.12.

**Table 2.** Mean saccade latency and amplitude of each condition in Experiment 2. Numbers in the parentheses are standard deviations (SDs).

|  | Upper visual field |  |  | Lower visual field |  |  |
| --- | --- | --- | --- | --- | --- | --- |
|  | Ecc. = 3° | Ecc. = 6° | Ecc. = 12° | Ecc. = 3° | Ecc. = 6° | Ecc. = 12° |
| *Latency (ms)* |  |  |  |  |  |  |
| Dist. to Hor. = 30° |  |  |  |  |  |  |
| Gap | 141<br>(11.82) | 129<br>(11.53) | 137<br>(9.95) | 150<br>(15.20) | 145<br>(11.82) | 140<br>(8.76) |
| Overlap | 185<br>(17.95) | 172<br>(21.88) | 191<br>(20.06) | 197<br>(12.10) | 201<br>(16.83) | 204<br>(22.22) |
| Dist. to Hor. = 75° |  |  |  |  |  |  |
| Gap | 148<br>(21.74) | 140<br>(16.06) | 146<br>(8.56) | 177<br>(19.22) | 169<br>(13.45) | 168<br>(8.91) |
| Overlap | 184<br>(25.01) | 179<br>(23.55) | 200<br>(8.39) | 233<br>(9.98) | 230<br>(11.14) | 235<br>(15.49) |
| *Amplitude (deg.)* |  |  |  |  |  |  |
| Dist. to Hor. = 30° |  |  |  |  |  |  |
| Gap | 2.54<br>(0.22) | 5.32<br>(0.34) | 10.34<br>(1.22) | 2.94<br>(0.37) | 5.60<br>(0.89) | 10.70<br>(1.15) |
| Overlap | 2.69<br>(0.22) | 5.58<br>(0.22) | 11.26<br>(0.93) | 2.98<br>(0.32) | 5.89<br>(0.62) | 11.33<br>(0.72) |
| Dist. to Hor. = 75° |  |  |  |  |  |  |
| Gap | 2.48<br>(0.36) | 4.71<br>(0.87) | 9.09<br>(1.39) | 2.94<br>(0.39) | 5.60<br>(0.60) | 10.70<br>(0.96) |
| Overlap | 2.52<br>(0.40) | 5.03<br>(0.89) | 10.16<br>(1.14) | 2.98<br>(0.33) | 5.89<br>(0.64) | 11.33<br>(1.17) |

**Figure 5.**
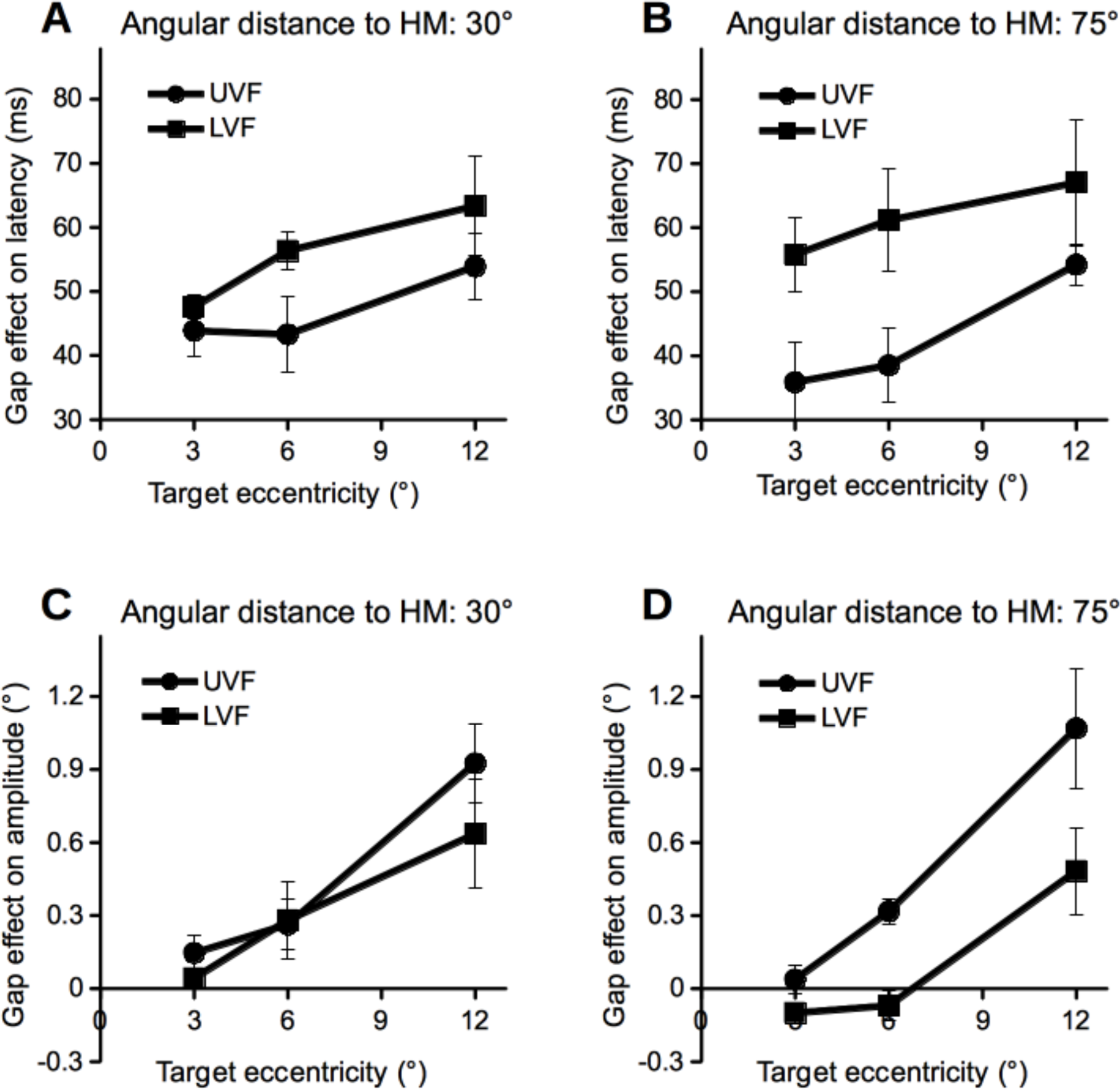
The effect of temporal gap on saccade latency and saccade amplitude. Error bars denote ±1 SEM.

### The temporal gap effect on saccade amplitude

As clearly shown in Table 2, we replicated previous findings (Wang & Theeuwes, 2014) insofar as saccades had larger amplitudes in the overlap condition (M = 6.48°, SD = 0.44) compared to the gap condition (M = 6.14°, SD = 0.50), *t*(3) = 8.93, *p* < 0.01. As noted above, the primary dependent measure of interest was the gap effect on saccade amplitude, i.e., the amplitude difference between overlap and gap conditions (see Figure 5C-D). An ANOVA on the gap effect revealed a significant effect of target eccentricity, *F*(1,3) = 34.45, *p* < 0.01; in the eccentricity range tested in the present experiment, the more eccentric the saccade target, the stronger the gap effect. With this critical observation one can reason that the stronger the gap effect, the more eccentric a sacccade target is *in colliculuar space*. Of prime interest was whether the gap effect was the same for UVF and LVF targets presented at the same eccentricity. If the representation of eccentricities in the LVF is compressed, then in collicular space LVF saccade targets will be less eccentric than UVF saccade targets presented at the same eccentricity. Consequently, the gap effect should be weaker for the LVF. Confirming this prediction, the main effect of visual field reached marginal significance, *F*(1,3) = 8.58, *p* = 0.061. As clearly shown in Figure 5C-D, the gap effect was much stronger for targets that appeared in the UVF. This effect was more prominent when the saccade target was 75° away from the horizontal meridian. All other effects were not significant, all *F* < 3.6, all *p* > 0.16.

## Discussion

The purpose of Experiment 2 was to examine whether the eccentricities in the UVF and LVF are mapped symmetrically in the SC motor map, taking advantage of the fact that the manipulation of temporal gap/overlap modulates saccade amplitudes. As clearly shown in Figure 5C-D, the gap effect on saccade amplitude generally increased with target eccentricity. One may argue that because more eccentric visual space is represented by increasingly smaller collicular area and thus, this eccentricity effect may have been confounded by the collicular size of the saccade target, which reduced as eccentricity increased. This is a reasonable argument, however, it did not make the methodology of Experiment 2 less-than-optimal. First, increasing the size of more eccentric saccade targets may reduce saccade accuracy and thus introduce a serious methodological confound. Second, previous cell-recording studies clearly showed that the strength of SC neuronal activation is relatively unaffected by the size of the saccade target, as long as it is smaller than the response field of the SC neuron (Schiller & Stryker, 1972). Also, the dependent measure of primary interest was the amplitude difference between gap and overlap conditions. This gap effect should be modulated by eccentricity similarly across visual fields, even if there was a confound with eccentricity. This was *not* what we observed in Experiment 2.

The observation that the gap effect increases with saccade target eccentricity provides us with a critical tool we can use to infer the relative target eccentricity in collicular space. That is, because the amplitude of the gap effect increases with target eccentricity, it stands to reason that the stronger the gap effect, the more eccentric a saccade target should be in colliculuar space. As shown in Figure 5D, the gap effect was much stronger in the UVF when the saccade target was relatively far from the horizontal meridian. We suggest that one can infer from this finding that saccade targets in the UVF are mapped to more caudal sites in the SC, which would be true if the representation of eccentricities is relatively compressed in the LVF.

## General discussion

Previous work has shown that a) the presence of visual distractors modulates the direction of saccades, and b) a temporal gap between fixation offset and saccade target onset will reduce the amplitude of saccades. Taking advantage of these two well-established behavioral effects, two experiments were carried out to examine whether the UVF and LVF are symmetrically represented in oculomotor maps, as assumed in widely accepted mathematical models of the SC (Ottes, Van Gisbergen, & Eggermont, 1986). Experiment 1 found that, when the saccade target and a visual distractor were presented at locations mirroring the horizontal meridian, the directional deviation caused by the distractor was much stronger for LVF targets. Experiment 2 found that the temporal gap between the offset of the fixation stimulus and the onset of the saccade target modulated the amplitude of saccades and this modulation was weaker for saccades towards LVF targets. We suggest that these results are consistent with the notion that the representation of both meridians and eccentricities in the LVF are compressed in oculomotor maps (see the Discussion sections of Experiments 1 & 2). We suggest that this proposition deserves further neurophysiological explorations for two reasons.

First, anatomical asymmetries have been shown in various retinotopic brain maps. For instance, Van Essen et al. (1984) showed that the striate area devoted to the peripheral part of the LVF was, on average, about 1.25 times of that devoted to the UVF. A ratio of 1.22 was later reported for the central visual field by Tootell et al. (1988). A similar asymmetry also exists in the LGN, with a ratio of 1.13 (Connolly & Van Essen, 1984). In an attempt to map the visual projections to the mouse superior colliculus, Drager and Hubel (1976) found that the UVF is disproportionally represented. The magnification factor was higher for regions 30° above the horizontal meridian, suggesting that a larger tectal surface is devoted to a point in the UVF than in the LVF. This observation is consistent with our claim that the LVF is underrepresented in the oculomotor map in the SC.

Second, a compressed representation of the LVF in oculomotor maps provides a straight-forward and consistent explanation for several important behavioral findings. For example, if the representation of eccentricities in the LVF is relatively compressed, neuronal activations related to saccades into the LVF should be closer to the rostral pole of the SC and thus, suffer greater suppression. As a result, saccades into the LVF should take longer to initiate. This is what researchers found in previous work (e.g., Abegg et al., 2015) and in the present experiments. Also, as in our Experiment 1, previous work has shown that the magnitude of trajectory deviations caused by distractors is stronger for targets in LVF (Petrova, Wentura, & Fu, 2013). The simplest explanation is that, even if the deviation is comparable across the UVF and LVF in collicular space, because the representation of saccade directions is compressed in the LVF, the same collicular deviation for LVF saccades actually corresponds to a larger change in saccade direction.

The mapping between the visual field and the striate cortex is described with a complex logarithmic model (Schwartz, 1977). Mathematically equivalent mapping functions have been suggested by Ottes et al. (1986) to describe the mapping of the visual field in the SC. What these functions essentially do is to transform the polar coordinates of the visual field into the Cartesian coordinates used to describe the anatomical structure of the SC. This model contains two formulae (Eq. 1 & 2). In these formulae, *u* represents the anatomical distance (in mm) from the rostral pole in the SC along the axis representing the horizontal position (see Figure 1B), *v* is the perpendicular distance, and R and Φ are retinal eccentricity and the polar direction of the saccade target (in degrees). By approximating the the SC motor map constructed by Robinson (1972), Ottes et al. (1986) estimated that *B_u_* = 1.4 mm, *B_v_* = 1.8 mm/rad, and A = 3°. *B_u_* and *B_v_* are scaling constants determining the size of the SC motor map along its *u* and *v* axes, respectively. As shown in the present study, the representation of polar directions and eccentricities in the LVF is compressed relative to that in the UVF. This LVF compression can be easily accommodated in the equations, by making *B_v_* or/and *B_u_* contingent on the polar direction of the saccade target. For the SC motor map illustrated in Figure 3A, in which the representation of meridians in the LVF is compressed, *B_v_* was scaled by a sigmoidal transformation of the polar direction (Φ) (Eq. 3). The transformation of the polar direction was largely linear (*β* = 0.2).

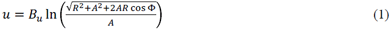

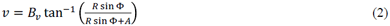

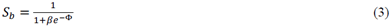

To conclude, the present results suggest that the LVF is underrepresented in oculomotor maps (e.g., the SC). Previc (1991) has argued that the near (peripersonal) and far (extrapersonal) spaces roughly correspond to the LVF and UVF, respectively. The oculomotor system, which plays a critical role visual search and orientation, is biased towards the processing of information in the UVF (far space). This functional bias may be the shaping force of the disproportional representation of the UVF in oculomotor maps.

